# Phase separation of both a plant virus movement protein and cellular factors support virus-host interactions

**DOI:** 10.1101/2021.05.11.443547

**Authors:** Shelby L. Brown, Jared P. May

## Abstract

Phase separation concentrates biomolecules, which should benefit RNA viruses that must sequester viral and host factors during an infection. Here, the p26 movement protein from *Pea enation mosaic virus 2* (PEMV2) was found to phase separate and partition in nucleoli and G3BP stress granules (SGs) *in vivo*. Electrostatic interactions drive p26 phase separation as mutation of basic (R/K-G) or acidic (D/E-G) residues either blocked or reduced phase separation, respectively. During infection, p26 must partition inside the nucleolus and interact with fibrillarin (Fib2) as a pre-requisite for systemic trafficking of viral RNAs. Partitioning of p26 in pre-formed Fib2 droplets was dependent on p26 phase separation suggesting that phase separation of viral movement proteins supports nucleolar partitioning and virus movement. Furthermore, viral ribonucleoprotein complexes containing p26, Fib2, and PEMV2 RNA were formed via phase separation *in vitro* and could provide the basis for self-assembly *in planta*. Interestingly, both R/K-G and D/E-G p26 mutants failed to support systemic trafficking of a *Tobacco mosaic virus* (TMV) vector in *Nicotiana benthamiana* suggesting that p26 phase separation, proper nucleolar partitioning, and systemic movement are intertwined. p26 also partitioned in SGs and G3BP over-expression restricted PEMV2 accumulation >20-fold. Expression of phase separation-deficient G3BP only restricted PEMV2 5-fold, demonstrating that G3BP phase separation is critical for maximum antiviral activity.

**AUTHOR SUMMARY:** Phase separation of several cellular proteins is associated with forming pathological aggregates and exacerbating neurodegenerative disease progression. In contrast, roles for viral protein phase separation in RNA virus lifecycles are less understood. Here, we demonstrate that the p26 movement protein from *Pea enation mosaic virus 2* phase separates and partitions with phase-separated cellular proteins fibrillarin and G3BP. The related orthologue from *Groundnut rosette virus* has been extensively studied and is known to interact with fibrillarin in the nucleolus as a pre-requisite for virus movement. We determined that basic residues and electrostatic interactions were critical for p26 phase separation. Furthermore, mutation of charged residues prevented the rescue of a movement-deficient *Tobacco mosaic virus* vector in *Nicotiana benthamiana*. Stress granules form through phase separation and we found that p26 could partition inside stress granules following heat shock. Phase separation of the stress granule nucleator G3BP was required for maximum antiviral activity and constitutes a host response that is dependent on cellular protein phase separation. Collectively, we demonstrate that phase separation of a plant virus protein facilitates virus-host interactions that are required for virus movement and phase separation of cellular proteins can simultaneously restrict virus replication.

## INTRODUCTION

Cellular organelles are membrane-bound compartments that are critical for eukaryotic cell function and RNA viruses often co-opt organelles to promote virus replication. Organelles exploited by RNA viruses include the endoplasmic reticulum (ER) [1], mitochondria [2], nucleus [3], and Golgi apparatus [4]. Recently, much attention has been directed towards membraneless organelles that form through protein phase separation. Phase separation transforms a single-phase solution into a dilute phase and droplet phase that concentrates biomolecules, such as proteins or RNAs [5, 6]. Some cellular proteins phase separate and form aggregates that are associated with several neurodegenerative disorders [7]. Proteins that undergo phase separation consistently contain intrinsically disordered regions (IDRs) that self-associate to form oligomers [8]. Many IDR-containing proteins have RNA-recognition motifs that non-specifically bind RNA and fine-tune phase separation by controlling material exchange, shape, and rigidity of liquid droplets [8, 9]. Proteins that phase separate are often enriched in arginine residues that can participate in cation-pi interactions with aromatic contacts and promote phase separation [10].

Membraneless organelles exist as liquids, gels, or solids, [11]. The most notable examples of liquid-liquid phase separated (LLPS) membraneless compartments are the nucleolus and P-bodies in the cytoplasm [12]. Less dynamic stress granules (SGs) also form in the cytoplasm through phase separation and allow host cells to repress translation and influence messenger RNA (mRNA) stability in response to various stresses [13]. SGs are visible by microscopy within minutes following stress and contain Ras-GTPase-activating protein SH3 domain-binding protein 1 (G3BP1) that self-associates to induce SG formation [14]. SGs contain a stable inner core and an outer shell that is formed by weak electrostatic and/or hydrophobic interactions [15]. The G3BP1 inner core is resistant to dilution (atypical for LLPS) and has been considered to be a form of liquid-solid demixing [16]. Interestingly, G3BP1 can have either proviral [17–19] and antiviral roles [20–22] in RNA virus lifecycles.

Members of the *Mononegavirales*, including *Rabies virus, Measles virus* (MeV), and *Vesicular stomatitis virus* generate phase-separated cytoplasmic inclusion bodies that create viral factories [23–25]. Phase separation of MeV N and P proteins also promotes efficient encapsidation of viral RNAs [25]. Several groups have recently demonstrated that the nucleocapsid (N) protein from the novel SARS-CoV-2 coronavirus undergoes LLPS [26]. SARS-CoV-2 N protein phase separation is stimulated by the 5’ end of its cognate RNA [27] and can partition into phase separations of heterogeneous nuclear ribonucleoproteins like TDP-43, FUS, and hnRNPA2 [28]. The SARS-CoV-2 N protein also interacts with G3BP1 and can attenuate SG formation [29, 30].

*Pea enation mosaic virus 2* (PEMV2) is a small (4,252 nt), positive-sense RNA plant virus in the tombusvirus family. The PEMV2 long-distance movement protein (MP) p26 is required for systemic trafficking of viral RNA throughout an infected plant. Both p26 and the orthologue pORF3 from *Groundnut rosette virus* (GRV) primarily localize to the cytoplasm, but also target cajal bodies in the nucleus and eventually partition in the nucleolus [31–33]. Umbravirus ORF3 proteins must interact with nucleolar fibrillarin (Fib2), a pre-requisite for long-distance movement of viral RNA [33–35]. Additionally, the polerovirus *Potato leafroll virus* (PLRV) and the potexvirus *Bamboo mosaic virus* satellite RNA (satBaMV) encode proteins that must also localize to the nucleolus and interact with fibrillarin to support systemic movement [36–38]. Fibrillarin phase separates and forms the dense fibrillar component (DFC) of the nucleolus that shares a similar structure to SGs [15, 39]. Although the nucleolus itself is a phase separation and several plant virus proteins co-localize with fibrillarin, the role of viral protein phase separation in plant virus lifecycles has not been investigated.

This study demonstrates that PEMV2 p26 undergoes phase separation both *in vitro* and *in vivo* and forms highly viscous condensates. Viral ribonucleoprotein (vRNP) complexes containing p26, Fib2, and PEMV2 RNA were reconstituted *in vitro* through phase separation and likely represents the version of the *in vivo* event necessary for systemic trafficking. Mutating charged residues required for phase separation and proper nucleolar localization blocked the movement of a viral vector suggesting that phase separation and virus movement are intertwined. Finally, p26 phase separates *in vivo* with the SG nucleator, G3BP, which exhibits strong antiviral activity towards PEMV2. PEMV2 accumulation was largely restored during expression of a phase-separation deficient G3BP, demonstrating that phase separation of select cellular proteins aids host antiviral responses.

## RESULTS

### p26 forms poorly dynamic condensates *in vivo*

p26 and related umbravirus orthologues form large cytoplasmic inclusion bodies during infection [35, 40, 41]. To define the material properties of p26 inclusion bodies *in vivo*, we used fluorescence recovery after photobleaching (FRAP) [42]. p26 with a C-terminal green fluorescent protein (GFP) tag was expressed from a *Cauliflower mosaic virus* (CaMV) 35S promoter in *Nicotiana benthamiana* by agroinfiltration (Fig. 1A). Separately, free GFP was expressed from a 35S promoter and was evenly distributed throughout the cytoplasm and nucleus of the cell (i.e, outside of the large vacuole that comprises most of the cellular space) (Fig. 1B, Left). In contrast, p26:GFP formed large cytoplasmic inclusion bodies as previously observed (Fig. 1B, Right) [41]. Nearly 50% recovery of p26:GFP was observed by 30 seconds post-bleach (Fig. 1C) demonstrating that p26 inclusion bodies have measurable fluidity. However, p26:GFP failed to recover any further suggesting that p26 forms poorly dynamic condensates *in vivo*, similar to what has been observed for G3BP1 SG cores [16].

**Fig. 1.**
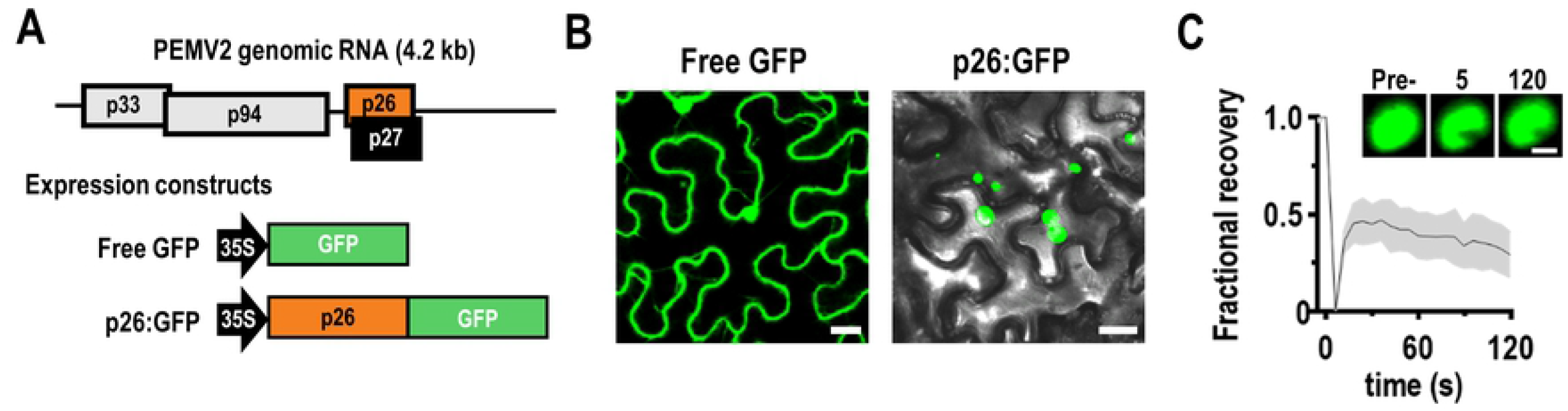
p26 forms poorly dynamic condensates *in vivo*. (A) PEMV2 is a small positive-sense RNA plant virus that encodes 4 genes, including the p26 long-distance movement protein. Free GFP and p26 C-terminally fused with GFP (p26:GFP) were expressed from binary expression plasmids under the constitutive CaMV 35S promoter (B) Following agroinfiltration of *N. benthamiana*, confocal microscopy showed diffuse cytoplasmic and nuclear expression of free GFP whereas p26:GFP formed large cytoplasmic bodies. Note that the majority of plant mesophyll cells is taken up by a single large vacuole. Differential interference contrast (DIC) microscopy was used for p26:GFP samples to visualize cell borders. Bar scale: 20 µm. (C) FRAP analysis of p26:GFP was performed by photobleaching cytoplasmic condensates and monitoring fluorescence recovery. A representative p26:GFP condensate is shown before photobleaching, immediately following photobleaching (5 s), and at 120 s. Bar scale 5 µm. Average FRAP intensity is shown from seven FRAP experiments and shaded area represents 95% confidence interval.

### p26 is intrinsically disordered and undergoes phase separation

To support the *in vivo* FRAP observations suggesting that p26 undergoes phase separation, *in vitro* assays were performed. Using the IUPred disorder prediction model [43], a large IDR spanning amino acids 1-132 was predicted in p26 (Fig. 2A). For comparison, the non-essential PEMV2 cell-to-cell movement protein, p27, did not contain a predicted IDR (Fig. 2A). Glycine, proline, and arginine amino acids are the most abundant residues in the p26 IDR (Fig. 2B), consistent with disordered proteins known to phase separate [44]. The p26 IDR was fused to the N-terminus of GFP and purified from *E. coli* for *in vitro* phase separation assays (Fig. 2C). 10% PEG-8000 was used to mimic cellular crowding and IDR-GFP readily phase separated under crowding conditions as observed by both turbidity assays (Fig. 2D) and confocal microscopy (Fig. 2E). In contrast, free GFP failed to phase separate under all tested conditions. High-salt concentrations disrupt self-associations resulting from electrostatic interactions and can reverse phase separation [45]. Accordingly, IDR-GFP concentrations near the saturation concentration (*C*_*sat*_ = 4 µM) failed to phase separate in the presence 800 mM NaCl and 1 M NaCl was required to block phase separation under standard assay conditions using 8 µM protein (Fig. 2E and F). IDR-GFP phase separations were next treated with 10% 1,6 hexanediol to probe the material properties of the *in vitro* condensates. 1,6 hexanediol interferes with weak hydrophobic protein-protein interactions and dissolves liquid-like, but not solid or highly viscous phase separations [46]. IDR-GFP phase separations were resistant to 1,6 hexanediol treatment (Fig. 2E) and FRAP analyses revealed that IDR-GFP condensates only reached 13% recovery after 2 minutes following photo-bleaching (Fig. 2J). Together, these data suggest that the p26 IDR drives phase separation through electrostatic interactions and the resulting condensates are highly viscous.

**Fig. 2.**
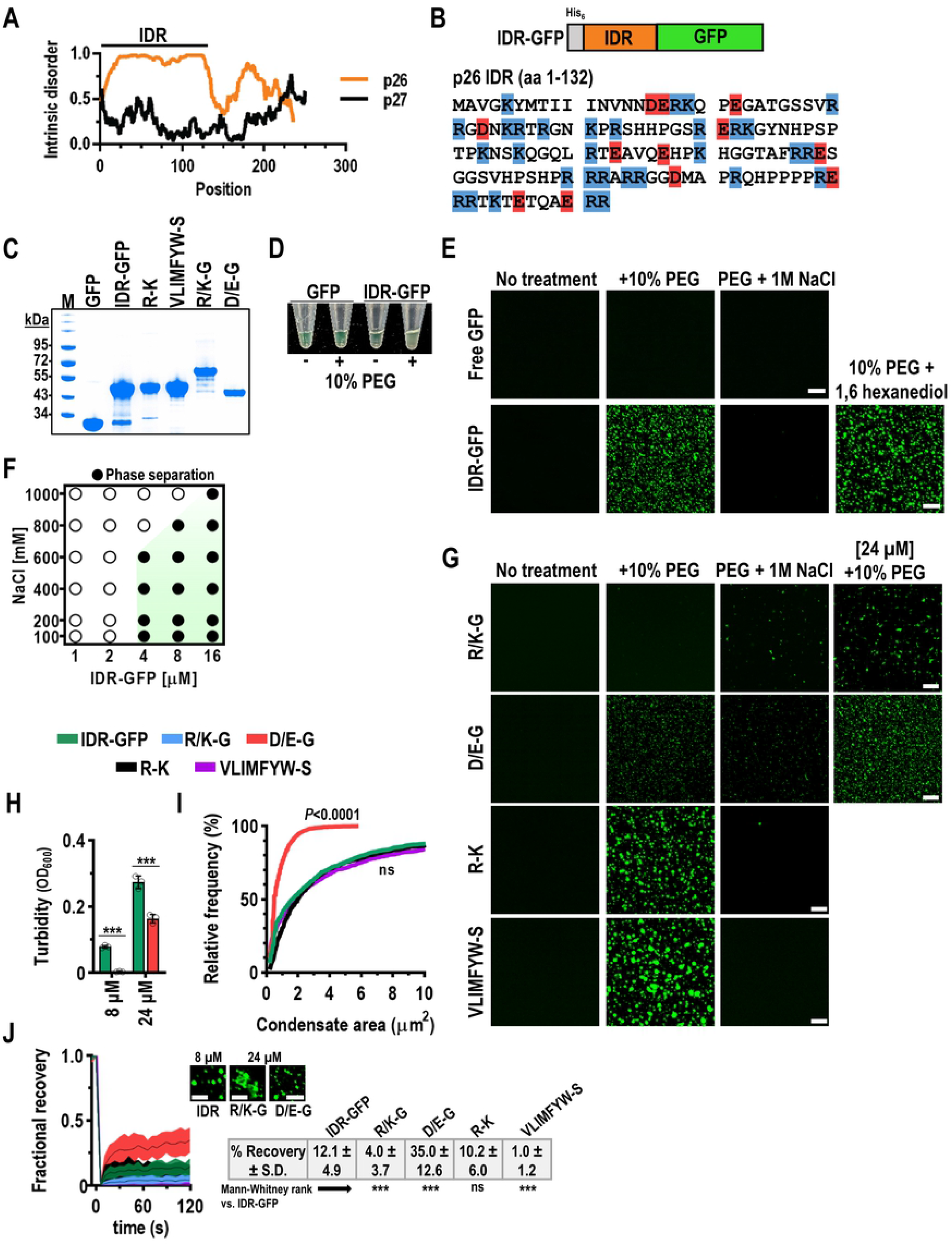
p26 is intrinsically disordered and phase separates through electrostatic interactions. (A) PEMV2 p26 contains a large intrinsically disordered region (IDR) spanning amino acids 1-132. The dispensable cell-to-cell movement protein, p27, is highly ordered. (B) The p26 IDR was fused to the N-terminus of GFP for bacterial expression and contained an N-terminal histidine tag. The p26 IDR sequence is shown with highlighted residues corresponding to basic (blue) or acidic (red) residues. (C) Recombinant proteins used in this study were analyzed by SDS-PAGE to assess size and purity. Proteins were stained using Coomassie Blue. Marker (M) sizes are shown in kilodaltons (kDa). Note: R/K-G ran markedly higher both *in vitro* and *in vivo* (see Fig. 4B). (D) Molecular crowding was induced with 10% PEG in the presence of 24 µM free GFP or IDR-GFP. The IDR-GFP solution became turbid in the presence of PEG, indicative of phase separation. (E) *In vitro* phase separation assays were visualized by confocal microscopy. 8 µM protein was used for all assays and 10% PEG-8000 was added as a crowding agent (Middle panels). One molar NaCl was added to disrupt electrostatic interactions (Right panel). 10% 1,6 hexanediol was added to IDR-GFP phase separations to assess the fluidity of condensates. Bar scale: 20 µm. (F) Phase diagram for IDR-GFP gives an apparent *C*_*sat*_ = 4 µM and sensitivity to high NaCl concentrations. Results are representative of two independent experiments. (G) IDR mutants (8 µM) were examined using *in vitro* phase separation assays. R/K-G formed irregular aggregates at high concentration (24 µM) and D/E-G showed reduced phase separation compared to IDR-GFP. R-K and VLIMFYW-S mutants appeared like wild-type IDR. Bar scale: 20 µm (H) D/E-G had significantly reduced turbidity (OD_600_) under crowding conditions when compared to IDR-GFP at 8 µM and 24 µM concentrations. Data represents three independent replicates for each condition. Bars denote standard deviations. *** *P<*0.001 unpaired t test (I) Mean condensate sizes for all mutants (excluding R/K-G) were plotted by cumulative distribution frequency. Particle sizes were measured from three representative 20x fields using ImageJ. *P* values represent results from two-tailed Mann-Whitney tests compared to IDR-GFP. ns: not significant. (J) FRAP was performed for *in vitro* condensates. 24 µM protein was used for R/K-G and D/E-G. Inset shows representative IDR-GFP and D/E-G droplets, or R/K-G aggregates. Bar scale: 10 µm. Table shows %recovery after 2 minutes with Mann-Whitney rank test comparisons against IDR-GFP. Data represents 7-10 separate FRAP measurements for each mutant. Shaded areas represent 95% confidence intervals.

### Charged residues are critical for efficient p26 IDR phase separation

To determine if specific groups of amino acids contribute to p26 phase separation, a series of IDR-GFP mutants were purified (Fig. 2C) and tested. First, all basic or acidic residues were mutated to glycine (R/K-G or D/E-G, respectively). Since high-salt blocks IDR-GFP phase separation, simultaneous mutation of either basic or acidic residues was predicted to inhibit phase separation. Indeed, R/K-G failed to phase separate while D/E-G showed significantly reduced phase separation compared to IDR-GFP when examined by confocal microscopy (Fig. 2G), turbidity assays (Fig. 2H), or mean condensate size (Fig. 2I). At elevated concentrations (24 µM), R/K-G formed non-uniform aggregates and failed to recover in FRAP assays (Fig. 2J). However, D/E-G condensates displayed significantly elevated fluidity when compared to IDR-GFP with 35% recovery after 2 minutes (Fig. 2J) and may be due to increased glycine content that has been associated with increasing condensate fluidity [47]. Cation-pi interactions between arginines and aromatic rings promote phase separation and are useful for predicting the propensity of a protein to phase separate [10, 48]. However, the p26 IDR only contains three aromatic residues that could potentially facilitate cation-pi interactions and mutation of all arginines to lysine (R-K) had no effect on phase separation, condensate size, or FRAP recovery (Fig. 2G-J). Finally, hydrophobic IDR residues (V, L, I, M, F, Y, W) were mutated to polar serine residues to reduce the hydrophobicity and prevent hydrophobic interactions that can drive phase separation [49]. Again, VLIMFYW-S phase separated like wild-type and was sensitive to high-salt (Fig. 2G). However, VLIMFYW-S condensates failed to recover in FRAP assays (Fig. 2J). These results suggest that hydrophobic residues contribute to the limited fluidity of p26 phase separations or rather the observed decrease in fluidity is due to the hardening properties of introduced serine residues [47].

### p26 partitions in the nucleolus and forms assemblies with the fibrillarin GAR domain via phase separation

Umbravirus movement proteins must access the nucleolus to support systemic virus trafficking [33]. Here, the nucleolar partitioning of wild-type or mutant p26:GFP was examined after agroinfiltration of *N. benthamiana* leaves. As previously reported for related orthologues [33–35, 50], p26 was observed in the nucleolus and cajal bodies in addition to forming cytoplasmic granules (Fig. 3A, Left). However, R/K-G p26 was diffusely expressed throughout the cytoplasm and failed to partition in the nucleolus (Fig. 3A, Middle). Conserved arginines in the related GRV pORF3 were previously shown to constitute a nuclear localization signal (NLS) [50]. Therefore, both p26 nuclear localization and phase separation are controlled by arginine residues and based on our mutagenesis studies it is unlikely that phase separation can be abolished without disrupting the NLS. Despite having markedly reduced phase separation *in vitro*, D/E-G p26 localized to the nucleolus and formed cytoplasmic granules that appeared like wild-type (Fig. 3A, Right). However, D/E-G had increased nucleolar retention compared to wild-type p26 as determined using the Manders Overlap Coefficient (MOC) to measure the degree of spatial overlap between D/E-G and DAPI-stained nuclei (Fig. 3B). Nucleolar localization/retention of *Arabidopsis thaliana* ribosomal proteins is dependent on the overall positive (basic) charge of the protein [51] and could explain the increased retention of D/E-G since the net charge of D/E-G at pH 7.4 is +36 compared to +14 for wild-type p26. Similarly, nucleolar accumulation of the *Human immunodeficiency virus 1* Tat protein strongly correlates with the overall net charge [52]. Together, these data demonstrate that basic residues are required for p26 nucleolar partitioning and the overall net charge influences nucleolar trafficking.

**Fig. 3.**
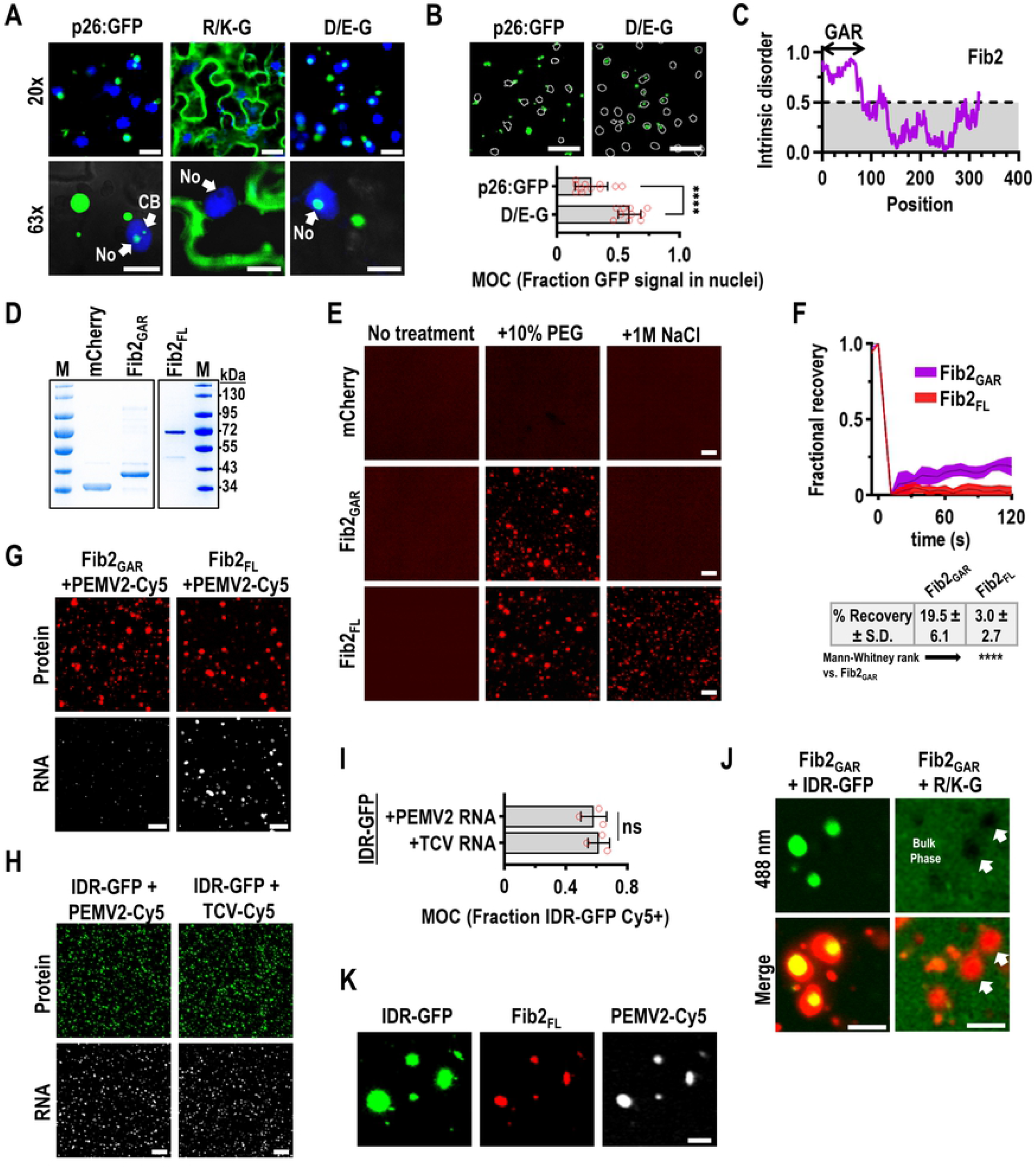
Phase separation supports p26 partitioning in Fib2 droplets and vRNP formation. (A) p26:GFP, R/K-G, and D/E-G GFP fusions were expressed in *N. benthamiana* leaves following agroinfiltration. Prior to imaging, leaves were infiltrated with 5 µg/mL DAPI to stain nuclei. 20x and 63x fields are shown. Arrows denote the nucleolus (No) or cajal bodies (CB). Bar scale: Top 20 µm; Bottom 10 µm. (B) Nuclear localization of p26:GFP or D/E-G was quantified using Mander’s overlap coefficient (MOC) using ImageJ and EzColocalization [80]. White outlines represent thresholded nuclei. Representative results are from ten 20x fields. Bar scale: 50 µm. Error bars denote standard deviations. \*\*\*\**P*<0.0001 unpaired t test. (C) Fib2 contains an N-terminal glycine- and arginine-rich (GAR) domain that is intrinsically disordered. (D) Either the Fib2 GAR domain (Fib2_GAR_) or full-length Fib2 (Fib2_FL_) were fused to mCherry and purified from *E. coli* and analyzed by SDS-PAGE. Molecular weight (kDa) marker is shown. (E) mCherry, Fib2_GAR_, and Fib2_FL_ were examined by confocal microscopy after inducing phase separation with 10% PEG-8000 alone or in the presence of 1 M NaCl. 8 µM protein was used for all assays. Bar scale: 20 µm. (F) FRAP analyses of Fib2_GAR_ and Fib2_FL_ condensates. Shaded areas represent 95% confidence intervals. Results are from 8 separate FRAP experiments. Table shows %recovery after two minutes. **** *P<*0.0001 Mann-Whitney rank test comparison Fib2_GAR_ and Fib2_FL_ droplets were pre-formed prior to addition of PEMV2-Cy5 at a 500:1 protein:RNA molar ratio. PEMV2 RNA was only sorted to Fib2_FL_ condensates. Bar scale: 20 µm. IDR-GFP droplets were pre-formed prior to addition of PEMV2-Cy5 or TCV-Cy5 at a 500:1 protein:RNA molar ratio. Bar scale: 20 µm. (I) The fraction of IDR-GFP signal that was positive for Cy5-labelled RNA was determined by MOC analysis using EzColocalization [80]. ns: not significant by unpaired t test. Bars denote standard deviations. Three 20x fields were quantified for each condition. (J) Fib2_GAR_ droplets were pre-formed using 24 µM protein before the addition of 4 µM IDR-GFP or R/K-G. Sorting of IDR-GFP to Fib2 droplets was observed whereas R/K-G remained in the bulk phase and failed to partition in Fib2_GAR_ droplets. Bar scale 10 µm. (K) IDR-GFP, Fib2_FL_, and PEMV2-Cy5 RNA were mixed at a 500:500:1 molar ratio after pre-forming Fib2_FL_ and IDR-GFP condensates under crowding conditions. Droplets containing all components were observed. Bar scale: 10 µm. Images in all panels are representative of at least three independent experiments.

Fibrillarin (Fib2) is a known host factor required for systemic trafficking of umbravirus vRNPs [31, 32] and makes up the dense fibrillar component of the nucleolus [53]. The *A. thaliana* Fib2 N-terminus contains an intrinsically disordered glycine- and arginine-rich (GAR) domain (Fig. 3C) that is common to fibrillarin across eukaryotes [54]. To determine whether the GAR domain of *A. thaliana* Fib2 is sufficient for phase separation, the GAR domain (amino acids 7-77, Fib2_GAR_) was fused to the N-terminus of mCherry and purified from *E. coli* for *in vitro* phase separation assays (Fig. 3D). Full-length Fib2 was also fused to mCherry (Fib2_FL_) for comparison. Free mCherry did not phase separate in the presence of 10% PEG-8000 or under high-salt conditions (Fig. 3E). Fib2_GAR_ readily phase separated under crowding conditions but was unable to phase separate in the presence of 1 M NaCl (Fig. 3E). These results indicate that the GAR domain is sufficient to drive Fib2 phase separation through electrostatic interactions and is consistent with findings using mammalian or *Caenorhabditis elegans* fibrillarin [39, 55, 56]. Full-length Fib2 phase separated under crowding conditions but unlike Fib2_GAR_, Fib2_FL_ was resistant to 1 M NaCl (Fig. 3E). These results suggest that Fib2_FL_ condensates are not strictly dependent on electrostatic interactions or Fib2_FL_ forms aggregates that are resistant to high salt. Indeed, Fib2_FL_ condensates failed to recover in FRAP assays while Fib2_GAR_ droplets were poorly dynamic but recovered nearly 20% after two minutes (Fig. 3F). Earlier work has determined that the GAR domain increases the solid-like properties of fibrillarin condensates [55] and supports our observations that both Fib2_GAR_ and Fib2_FL_ are poorly dynamic.

### vRNPs required for systemic trafficking can be reconstituted *in vitro* via phase separation

Fib2 is a necessary component of umbravirus vRNPs that move systemically during infection. To determine whether full-length PEMV2 RNA could be sorted to Fib2 phase separations, Cy5-labelled PEMV2 RNA was mixed with pre-formed Fib2_GAR_ or Fib2_FL_ droplets at a 500:1 protein:RNA molar ratio. PEMV2-Cy5 RNA was not efficiently sorted into Fib2_GAR_ droplets (Fig. 3G) and is consistent with earlier findings that determined the GAR domain does not bind RNA [54, 55]. However, Fib2_FL_ efficiently captured PEMV2-Cy5 RNAs demonstrating that viral RNAs can partition with Fib2 phase separations (Fig. 3G). Since p26 must also bind PEMV2 RNA prior to trafficking, PEMV2-Cy5 RNA was mixed with pre-formed IDR-GFP droplets. Approximately 50% of IDR-GFP signal spatially overlapped PEMV2-Cy5 signal when visualized by confocal microscopy and quantified by MOC (Fig. 3H and I). Interestingly, partitioning of viral RNA inside IDR-GFP condensates was not unique to PEMV2 RNA since the distantly related *Turnip crinkle virus* (TCV) RNA was sorted to IDR-GFP phase separations with similar propensity as measured by MOC (Fig. 3H and I). Collectively, these results demonstrate that both cognate and non-cognate viral RNAs are readily sorted into p26 phase separations.

Since the related GRV pORF3 directly interacts with the Fib2 GAR domain [34], IDR-GFP was added to pre-formed Fib2_GAR_ droplets at a 1:6 molar ratio to determine whether p26 can partition into phase separated Fib2 condensates. Expectedly, IDR-GFP was readily sorted into pre-formed Fib2_GAR_ droplets *in vitro* (Fig. 3J, Left) and is likely the reconstituted version of the p26-Fib2 interaction required for Fib2 export from the nucleus and subsequent vRNA association. To determine whether phase separation of p26 supports Fib2 partitioning, the phase separation-deficient R/K-G mutant was added to pre-formed Fib2_GAR_ droplets. Interestingly, R/K-G remained in the bulk phase and was excluded from Fib2_GAR_ droplets (Fig. 3J, Right, White arrows) suggesting that the ability of p26 to phase separate supports the key interaction with Fib2 required for virus movement. Finally, Fib2_FL_ and IDR-GFP phase separation was induced by molecular crowding prior to the addition of PEMV2-Cy5 RNA. Droplets containing IDR-GFP, Fib2_FL_, and PEMV2 RNA were observed (Fig. 3K) and demonstrate that the critical p26-Fib2-RNA interaction necessary for systemic trafficking of PEMV2 RNAs can be reconstituted using *in vitro* phase separation assays. In summary, these findings support a role for p26 phase separation in supporting virus movement.

### Phase separation-deficient p26 mutants fail to systemically traffic a virus vector

To determine whether phase separation-deficient p26 mutants could support virus trafficking, a *Tobacco mosaic virus* vector was used to express free GFP, p26, R/K-G, or D/E-G GFP fusions (Fig. 4A). The TMV vector (pJL-TRBO) contains a coat protein (CP) deletion that has been previously reported to block systemic movement [57]. Interestingly, GRV pORF3 and PEMV2 p26 have been previously shown to systemically traffic TMV when expressed from a subgenomic promoter in place of CP [41, 58]. Local infections were established in young *N. benthamiana* plants (4^th^ leaf stage) and high levels of free GFP and lower levels of p26:GFP, R/K-G, and D/E-G were observed at 4 days post-infiltration (dpi) (Fig. 4B). Systemic trafficking of TMV:p26:GFP was readily apparent by 14 dpi by both visual inspection of leaves and RT-PCR (Fig. 4C). However, TMV expressing GFP, R/K-G, or D/E-G GFP fusions failed to move systemically at 14 dpi. Basic amino acids are known to function as a NLS for GRV pORF3 [50] and are also required for partitioning in pre-formed Fib2 droplets (Fig. 3J). Therefore, p26 nucleolar localization and phase separation are co-dependent on basic residues and the R/K-G mutation presumably blocks interactions with Fib2 and subsequent virus trafficking. Failure of D/E-G to support virus movement was surprising since D/E-G retained the ability to phase separate (albeit less efficiently) and localize to the nucleolus (Figs. 2G and 3A). However, increased nucleolar retention of D/E-G could contribute to the block in systemic movement and suggests that nucleolar and virus trafficking by p26 is a tightly regulated process. Together, these data suggest that p26 phase separation, nucleolar partitioning, and virus movement are connected and co-dependent on charged residues. The TMV CP deletion has been previously reported to block systemic movement of the TRBO vector [57], but we routinely observed systemic trafficking of pJL-GFP after 3 weeks (Supplemental Fig. 1). However, pJL-GFP was largely restricted to the petiole and midrib of systemic leaves whereas pJL-p26:GFP spread throughout the veins and invaded the lamina. Weak D/E-G GFP expression was observed in the petioles and midribs of upper leaves at 21 dpi while R/K-G GFP was not visible (Supplemental Fig. 1).

**Fig. 4.**
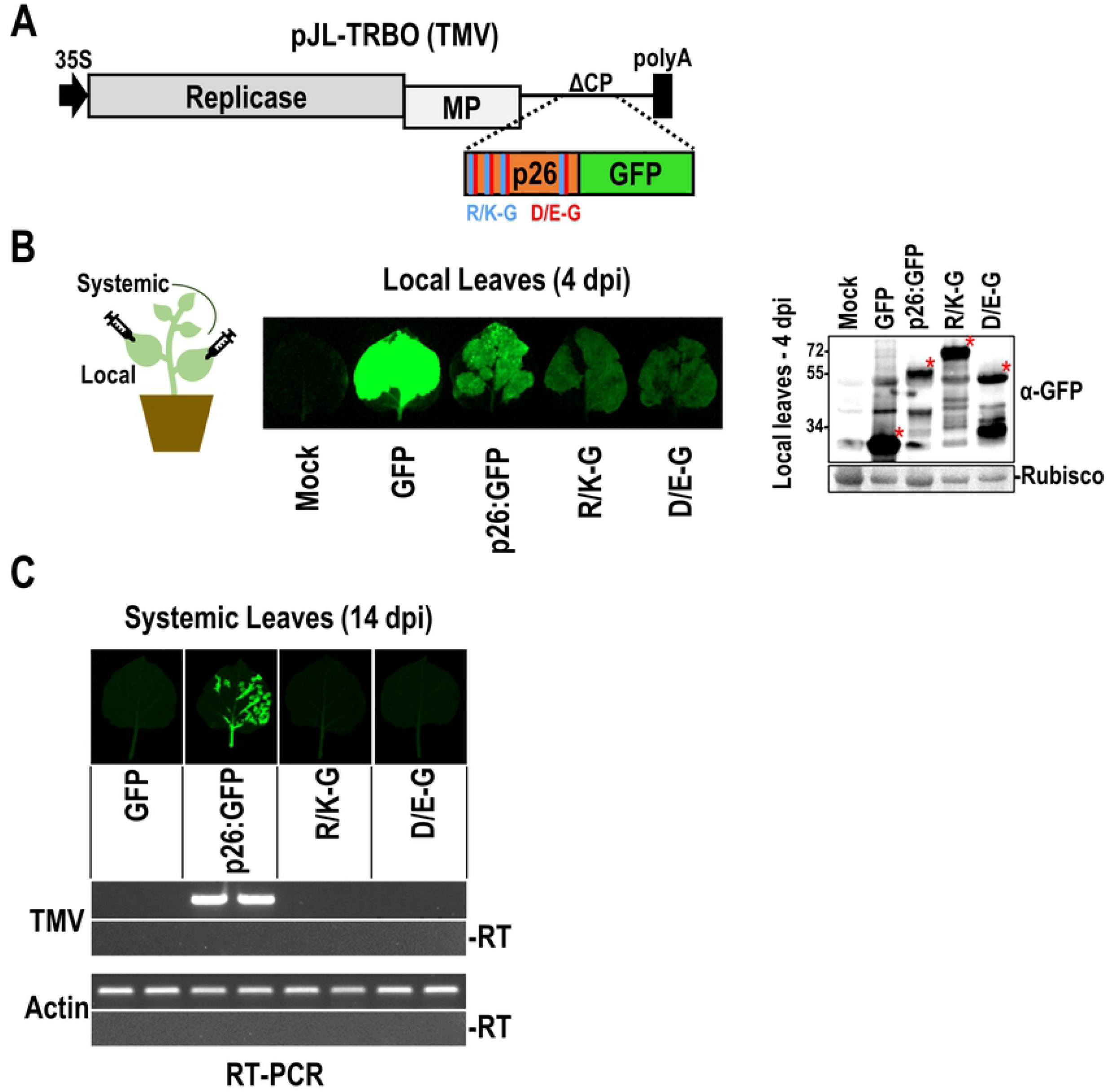
Phase separation-deficient p26 mutants fail to systemically traffic a virus vector. (A) pJL-TRBO TMV vector lacks CP and is severely impaired in systemic trafficking. Free GFP, p26:GFP, R/K-G, and D/E-G GFP fusions were inserted into pJL-TRBO to test whether systemic trafficking could be restored. (B) Following agroinfiltration of *N. benthamiana* leaves, TMV infections were established in local leaves. Free GFP, or GFP-fusion proteins were visualized and detected in local leaves at 4 dpi by UV exposure (Left) or western blotting (Right). Rubisco serves as a loading control. Red asterisks denote free GFP or GFP-fusion bands. (C) At 14 dpi, systemic leaves were imaged prior to total RNA extraction. RT-PCR was used to amplify 100-200 bp fragments targeting either the TMV replicase or actin as a control. - RT: No reverse transcriptase controls. Two pools of 3-4 leaves are shown for each construct. Results are representative of three independent experiments consisting of at least 4 plants/construct.

### p26 is sorted into G3BP phase separations that restrict PEMV2 accumulation

Our findings suggest that p26 phase separations share similar material properties to G3BP SG cores, mostly consistent with liquid-solid demixing [16]. A NTF2-RRM domain-containing protein from *A. thaliana* (AtG3BP) functions as a G3BP-like SG nucleator in plants [59]. The N-terminal NTF2 domain is required for both phase separation and recruitment to SGs [60, 61] and G3BP contains downstream IDRs (Fig. 5A). As previously demonstrated by Krapp et. al. [59], G3BP:RFP displays a diffuse cytoplasmic expression pattern under no stress, but forms cytoplasmic SGs after heat shock (Fig. 5B). As expected, ΔNTF2-G3BP failed to phase separate and form SGs following heat shock (Fig. 5B). When co-expressed with p26:GFP, recruitment of p26 to G3BP SGs was observed following heat shock (Fig. 5B). To determine if p26 partitions into SGs during a viral infection, G3BP:RFP was expressed in *N. benthamiana* plants systemically infected with TMV expressing p26:GFP (Fig. 5C). p26:GFP condensates co-localized with G3BP:RFP demonstrating that p26 and G3BP can share phase separations during an infection (Fig. 5C). To determine if G3BP expression is up- or down-regulated during PEMV2 infection, native G3BP gene expression was measured by RT-qPCR at 3 dpi in PEMV2-infected *N. benthamiana* (Fig. 5D). PEMV2 infection led to a 61% increase in G3BP expression (Fig. 5D) in accordance with previous RNA-seq analyses that showed a 2-fold increase in G3BP expression under similar conditions [41]. To determine if G3BP exerts a pro- or anti-viral effect on PEMV2 accumulation, G3BP:RFP was over-expressed alongside PEMV2. PEMV2 accumulation was reduced >20-fold during G3BP over-expression demonstrating that G3BP exerts strong antiviral activity towards PEMV2 (Fig. 5E). Virus accumulation was largely restored (only 5-fold inhibition) during overexpression of ΔNTF2-G3BP demonstrating that phase separation of G3BP is required for maximal antiviral activity (Fig. 5E). Together, these data demonstrate that p26 partitions inside G3BP SGs and G3BP phase separation facilitates an antiviral virus-host interaction.

**Fig. 5.**
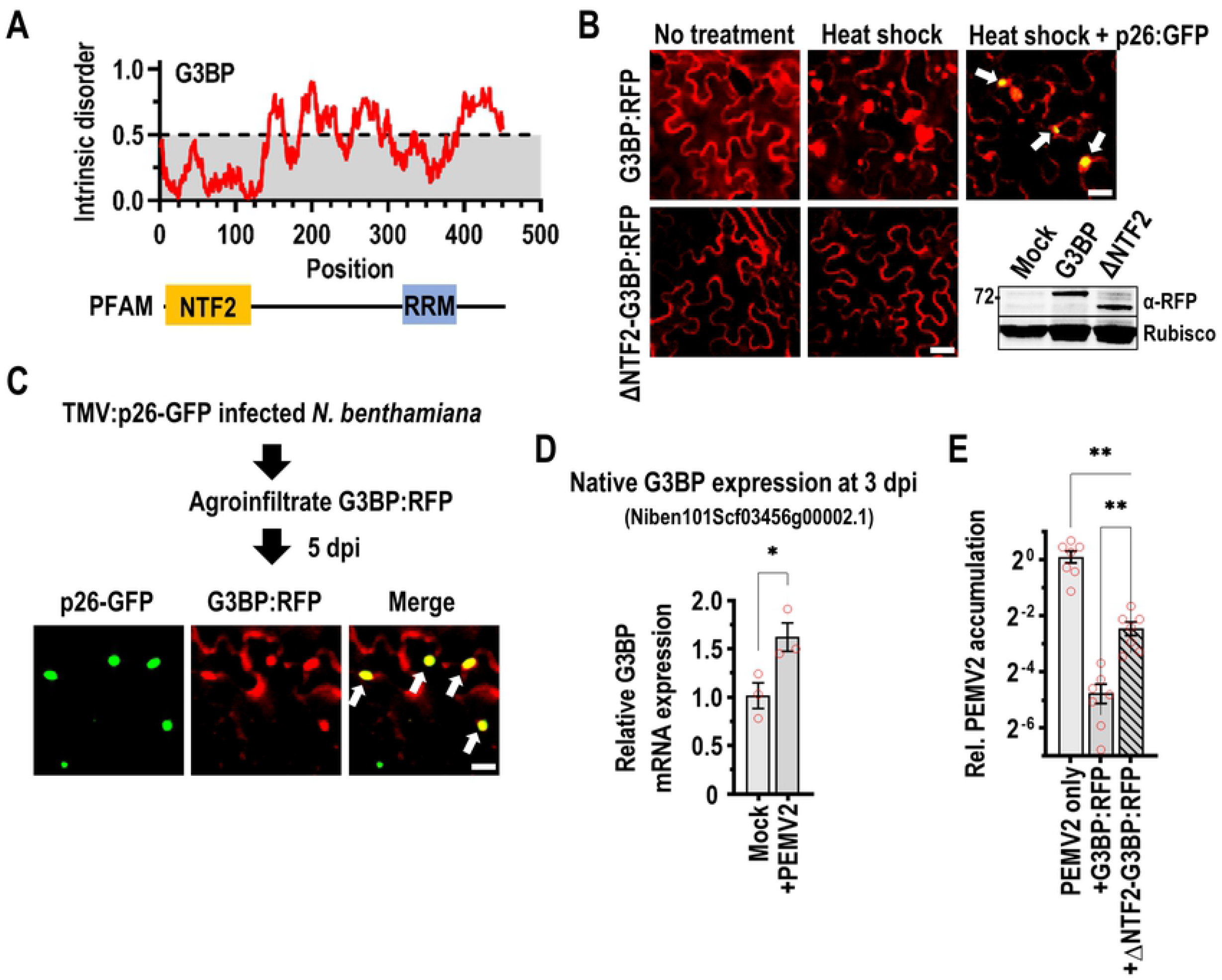
p26 is sorted into G3BP phase separations that restrict PEMV2 accumulation. (A) *A. thaliana* G3BP contains an ordered NTF2 domain and RNA recognition motif (RRM) in addition to intrinsically disordered regions. (B) G3BP:RFP or ΔNTF2-G3BP:RFP were agroinfiltrated into *N. benthamiana* leaves. At 3 dpi, plants were either imaged directly or heat shocked for 45 minutes at 37°C. p26:GFP was co-infiltrated with G3BP:RFP and p26 partitioning in G3BP SGs was observed (White arrows). Scale bar: 20 µm. Inset shows western blot using anti-RFP antibodies to detect full-length G3BP and ΔNTF2-G3BP. Rubisco was used as a loading control (C) G3BP:RFP was agroinfiltrated into systemically-infected TMV:p26:GFP plants to determine if p26 partitions in G3BP SGs during a virus infection. p26:GFP co-localized with G3BP SGs as labelled by white arrows. Scale bar: 20 µm. (D) Native G3BP expression was measured in Mock- or PEMV2-infected *N. benthamiana* by RT-qPCR. The co-agroinfiltrated p14 RNA silencing suppressor was used as a reference gene. Data is from three biological replicates. \**P*<0.05; student’s t-test. Bars denote standard error. (E) PEMV2 was agroinfiltrated alone, or alongside either G3BP or ΔNTF2-G3BP. At 3 dpi, total RNAs were extracted and used for RT-qPCR targeting PEMV2 or p14 (reference gene). Results shown are from 7 biological replicates from 2 independent experiments. Bars denote standard error. Brown-Forsythe and Welch ANOVA with multiple comparisons was used to determine if observed differences were significant. ** *P*<0.01.

## DISCUSSION

Phase separation of viral proteins has largely been associated with negative-sense RNA virus proteins that undergo phase separation to form virus factories [25], including Negri bodies during Rabies virus infections [23, 62, 63]. Also, measles virus N and P proteins encapsidate viral RNA more efficiently in a phase-separated droplet compared to a single phase solution [64]. In contrast, many positive-strand RNA viruses, including members of the *Tombusviridae* family form membranous replication organelles to concentrate virus replication complexes [65, 66]. While specific roles for phase separation of positive-sense RNA virus proteins in the virus lifecycle remain limited, phase separation of the SARS-CoV-2 N protein has been suggested to mediate nucleocapsid assembly and genome processing [67].

This study demonstrates that the p26 movement protein from the positive-sense RNA plant virus PEMV2 phase separates to form poorly dynamic condensates. Electrostatic interactions between acidic and basic IDR residues drive p26 phase separation and mutation of basic residues (R/K-G) abolished phase separation. Surprisingly, mutation of acidic residues (D/E-G) did not abolish phase separation but was significantly reduced compared to wild-type *in vitro*. Previous studies have found that phase separation of arginine-rich peptides can occur through charge repulsion in the presence of buffer counteranions and could support D/E-G phase separation [68, 69].

p26 must interact with fibrillarin (Fib2) in phase-separated nucleoli to support systemic virus trafficking [34], and conserved arginine residues have been shown to function as a NLS for the related GRV pORF3 [50]. Our results demonstrated that p26 nuclear localization and phase separation are both governed by basic amino acids making it problematic to separate these phenomena. However, the R/K-G IDR failed to accumulate in pre-formed Fib2_GAR_ droplets *in vitro* suggesting that phase separation of p26 could be required to partition in Fib2 phase separations and the nucleolus. Unsurprisingly, R/K-G p26 failed to support systemic movement of a TMV vector demonstrating that nucleolar partitioning, and potentially phase separation is required for virus movement. Mutation of acidic residues (D/E-G) significantly increased nucleolar retention of p26 and could be the result of increased protein net charge that is known to correlate with increased nucleolar retention [52]. Interestingly, D/E-G p26 failed to systemically traffic a TMV vector suggesting that the interplay between p26 nucleolar localization and virus movement is tightly regulated. In summary, charged amino acids play a critical role in p26 phase separation, nucleolar partitioning, and systemic virus movement.

Stress granules can support or restrict RNA virus replication and are assembled by the self-association and phase separation of G3BP [60, 61]. Seven *A. thaliana* G3BP-like candidates have been identified [70] and share an N-terminal NTF2 domain that is required for phase separation of mammalian G3BP1 [61]. In this study, the previously characterized AtG3BP-2 (AT5G43960) [59] was used to determine whether p26 could partition in G3BP stress granules. After heat shock, p26 readily partitioned inside G3BP SGs and both p26 and G3BP co-localized during virus infection. G3BP expression was upregulated during PEMV2 infection suggesting that G3BP could be expressed as part of a concerted host response to infection. G3BP over-expression severely restricted PEMV2 infection but was partially restored during expression of ΔNTF2-G3BP, demonstrating that phase separation of G3BP is necessary for maximum antiviral activity.

Since PEMV2 accumulation was not fully restored during ΔNTF2-G3BP expression, G3BP retains measurable antiviral activity in the dilute state. Human G3BP1 has been shown to bind and promote the degradation of mRNAs with structured 3’ untranslated regions (3’ UTRs) in conjunction with upframeshift 1 (Upf1) as part of the structure-mediated RNA decay (SRD) pathway [71]. PEMV2 contains a highly structured 3’ UTR [72] and like many RNA viruses is inhibited by Upf1 [73, 74]. Therefore, G3BP over-expression could enhance SRD targeting of PEMV2 RNAs. It remains unclear if p26 partitioning into G3BP SGs is beneficial or detrimental for PEMV2 replication. However, p26 was previously shown to disrupt the Upf1-dependent nonsense-mediated decay (NMD) pathway [41] and Upf1 is known to localize to G3BP1 SGs [75]. Partitioning of p26 into G3BP SGs could potentially interfere with Upf1- or G3BP-dependent RNA decay pathways.

In summary, our findings demonstrate that a plant virus movement protein phase separates and partitions inside cellular phase separations, namely the nucleolus and stress granules. Since nucleolar partitioning is required for virus trafficking and G3BP SG formation severely restricts PEMV2 replication, our findings highlight both beneficial and detrimental virus-host interactions mediated by phase separation.

## ACKNOWLEDGEMENTS

We would like to thank Dr. Björn Krenz (Leibniz Institut DSMZ, Brunswick, Germany) for the generous gifts of the G3BP:RFP construct. We would also like to thank Dr. Jonathan Dinman and Dr. Anne Simon (University of Maryland) for their thoughtful insight. We would also like to thank Dr. Anne Simon for critically reading this manuscript.

## AUTHOR CONTRIBUTIONS

Conceptualization, J.P.M; Methodology, S.B. and J.P.M; Investigation, S.B. and J.P.M; Writing – Original Draft, J.P.M.; Writing – Review & Editing, S.B. and J.P.M; Supervision, J.P.M.

## COMPETING INTERESTS

The authors declare no competing interests.

## MATERIALS & METHODS

### Construction of expression vectors

For C-terminal GFP fusion recombinant protein production in *E. coli*, pRSET his-eGFP [76] was used as a backbone and was a gift from Jeanne Stachowiak (Addgene plasmid # 113551). Wild-type IDR was PCR amplified from a full-length PEMV2 infectious clone, whereas R-K, VLIMFYW-S, R/K-G, and D/E-G were synthesized (Integrated DNA Technologies) as double stranded DNA fragments before used in restriction digests and ligation. All fragments except for R/K-G and D/E-G were cloned into the *BamH*I restriction site and sequenced for directionality and accuracy. R/K-G and D/E-G were cloned into pRSET his-eGFP using both the *Nhe*I and *BamH*I restriction sites and sequenced for accuracy.

Fibrillarin (Fib2) was first amplified from cDNA synthesized from *Arabidopsis thaliana* seedling total RNA using primers Forward 5’-GCAGCA<u>GCTAGC</u>ATGAGACCTCCTCTAACTGGAAGTGG-3’ and Reverse 5’-CTGCTGC<u>GGATCC</u>AGCAGCAGTAGCAGCCTTTGGCTTC-3’ where the underlined sequences denote the *Nhe*I and *BamH*I restriction sites used to introduce the PCR fragment into pRSET-his-mCherry [77], a gift from Jeanne Stachowiak (Addgene plasmid # 113552). The resulting construct is full-length Fib2 with a C-terminal mCherry fusion (Fib2_FL_). The Fib2 GAR domain was PCR amplified from Fib2_FL_, digested, and ligated into the *Nhe*I and *BamH*I restriction sites of pRSET-his-mCherry to generate Fib2_GAR_.

The *Tobacco mosaic virus* (TMV) expression vector pJL-TRBO has been previously described [57] and was a gift from John Lindbo (Addgene plasmid # 80082). The TMV vector containing p26:GFP has also been previously described [41]. R/K-G and D/E-G GFP-fusion inserts were commercially synthesized (Integrated DNA Technologies). TMV vectors expressing free GFP, R/K-G or D/E-G GFP fusions were constructed by cloning respective PCR fragments into the *Pac*I and *Not*I restriction sites in pJL-TRBO. p26:GFP, R/K-G, and D/E-G GFP fusions were also PCR amplified and cloned into pBIN61 using *BamH*I and *Sal*I restriction sites to transiently express p26-fusions downstream of the constitutive *Cauliflower mosaic virus* (CaMV) 35S promoter. G3BP:RFP was a generous gift from Dr. Björn Krenz and has been previously described [59]. To construct ΔNTF2-G3BP:RFP, G3BP-RFP was PCR amplified with amino acids 2-125 of G3BP omitted. PCR amplification introduced forward *BamH*I and reverse *Sal*I restriction sites for cloning into pBIN61S. All DNA constructs used in this study were sequenced for accuracy.

### Fluorescence recovery after photobleaching (FRAP)

A ∼2 µm diameter region was photobleached with 100% laser power with subsequent recovery measured at 5 s intervals. Background regions and unbleached reference condensates were recorded as controls. FRAP was performed using a Zeiss LSM 510 Meta confocal microscope with a 20X objective and Zen 2009 software. Data analysis was performed as previously described [78]. Briefly, background intensity was subtracted, intensities were normalized to set the first post-bleach value to zero and presented as a fraction of the pre-bleach fluorescence intensity.

### Protein expression and purification

Histidine-tagged recombinant proteins were expressed in BL21(DE3) *E. coli* (New England BioLabs) using autoinduction Luria-Bertani (LB) broth and purified using HisPur cobalt spin columns (Thermo Scientific). Proteins were purified under denaturing conditions according to the manufacturer’s protocol using 8 M urea. All equilibration, wash, and elution buffers contained 1 M NaCl to disrupt electrostatic interactions and restrict phase separation. Following elution of recombinant proteins from the cobalt resin, proteins were re-folded through dialysis in buffer containing 10 mM Tris-HCl (pH 7.0), 300 mM NaCl, 1 mM EDTA, 1 mM dithiothreitol, and 10% glycerol as previously done for the related pORF3 from *Groundnut rosette virus* [40]. Urea was removed in a stepwise fashion by using dialysis buffers containing 4 M Urea, 1 M Urea, or no Urea. Proteins were concentrated using centrifugal filters and concentrations were measured using the Bicinchoninic acid (BCA) protein assay kit (Millipore Sigma). Proteins were aliquoted and stored at −80°C.

### Phase separation assays

GFP- or mCherry-tagged proteins were used at a final concentration of 8 µM unless otherwise noted. Phase separation assays consisted of the following mixture: 8 µM protein, 10 mM Tris-HCl (pH 7.0), 1 mM DTT, 100 mM NaCl, and 10% PEG-8000 to mimic cellular crowding. Phase separation occurred rapidly and samples were directly loaded onto glass slides for confocal microscopy using a Zeiss LSM 510 Meta confocal microscope with a 20x objective and appropriate filters. High-salt conditions included NaCl at a final concentration of 1 M and “no treatment” did not include PEG-8000. Phase separation assays were performed at least twice across two protein preparations. Turbidity assays comparing IDR-GFP and D/E-G were performed with either 8 µM or 24 µM protein under standard assay conditions. 100 µL reactions were placed at room temperature for 15 minutes prior to measuring OD_600_ using a 96-well plate reader. Cy5-labelled PEMV2 or TCV RNA was synthesized by T7 run-off transcription using *Sma*I-linearized full-length infectious clones. Cy5-UTP (APExBIO) was added to *in vitro* transcription reactions according to the HiScribe T7 Quick High Yield RNA Synthesis Kit protocol (New England Biolabs). RNAs were included in phase separation assays at a final concentration of 16 nM (500:1 protein:RNA ratio).

### Agroinfiltration

Expression constructs were electroporated into *Agrobacterium tumerfaciens* (C58C1 strain). Liquid cultures were passaged in media containing 20 µM acetosyringone 1 day prior to infiltration. Overnight cultures were pelleted and resuspended in 10 mM MgCl_2_, 10 mM MES-K [pH 5.6], and 100 µM acetosyringone. Infiltration mixtures contained the p14 RNA silencing suppressor from *Pothos latent virus* [79] at a final OD_600_ of 0.2. pBIN-GFP constructs, TMV vectors, and G3BP:RFP constructs were infiltrated at a final OD_600_ of 0.4. The full-length PEMV2 expression construct has been previously described [73] and was agroinfiltrated at a final OD_600_ of 0.4. Visualization of nuclei in p26:GFP, R/K-G, or D/E-G-expressing plants was achieved by infiltrating a solution of 5 µg/mL DAPI (4′,6-diamidino-2-phenylindole) into leaves 45 minutes prior to imaging. Heat shock of G3BP-expressing plants was performed by placing plants at 37°C for 45 minutes prior to imaging. To visualize G3BP:RFP alongside p26:GFP during virus infection, young *N. benthamiana* plants (3-4 leaf stage) were first infiltrated with TMV:p26:GFP. After strong p26:GFP signal was observed in the systemic leaves (typically ∼2-3 weeks), G3BP:RFP was agroinfiltrated and imaged at 5 dpi using a Zeiss LSM 510 Meta confocal microscope with a 20x objective. Plants were grown in a humidity-controlled chamber at 24°C, 65% humidity, and 12-hour day/night schedule (200 µmol m^-2^s^-1^).

### TMV movement assay and RT-PCR

pJL-TRBO derived TMV vectors expressing GFP or p26-GFP fusions were agroinfiltrated into young *N. benthamiana* plants (3-4 true leaf stage). GFP fluorescence in local and systemic leaves was monitored daily. At 4 dpi, robust local infections were evident, and leaves were harvested by grinding in liquid nitrogen. Total protein was extracted by resuspending leaf tissue in 1X PBS supplemented with 3% β-mercaptoethanol and protease inhibitor cocktail (Thermo Scientific). Samples were mixed with 6X Laemmli SDS buffer, boiled, and separated by SDS-PAGE. A semi-dry transfer method was used to transfer proteins to nitrocellulose for western blotting using anti-GFP antibodies (Life technologies) at a 1:5000 dilution. Anti-rabbit IgG conjugated with horseradish peroxidase was used as a secondary antibody again at 1:5000 dilution. Blots were visualized using the Pierce enhanced chemiluminescence kit (Thermo Scientific). Systemic leaves were harvested at 14 dpi for total RNA extraction using Trizol. 100 ng total RNA digested with RQ1 DNase (Promega) served as template for reverse transcription using iScript supermix (Bio Rad). No reverse transcriptase controls (-RT) were Included for all sample and primer sets. 1 µL cDNA was used as template for 25 cycles of PCR using GoTaq polymerase (Promega) targeting the TMV replicase using forward primer 5’ CCGCGAATCTTATGTGGAAT 3’ and reverse primer 5’ TCCTCCAAGTGTTCCCAATC 3’. *N. benthamiana* actin was amplified by 31 cycles of PCR as a loading control with forward primer 5’ TCCTGATGGGCAAGTGATTAC 3’ and reverse primer 5’ TTGTATGTGGTCTCGTGGATTC 3’.

### RT-qPCR

Agroinfiltrated “spots” were cut from leaves and stored at −80°C. Samples were ground in liquid nitrogen and total RNA was extracted using the Quick-RNA Plant Kit (Zymo Research). An on-column DNase I step was added using RQ1 DNase (Promega). Total RNAs were used as templates for SYBR green-based one-step reverse-transcriptase quantitative PCR (RT-qPCR) using the NEB Luna One-Step RT-qPCR kit (New England Biolabs). All primers were validated by standard curve analysis and had PCR efficiencies ranging from 90-110%. Native *N. benthamiana* G3BP (Transcript ID: Niben101Scf03456g00002.1) was targeted using primers Forward 5’ TAGGGGAAGCAATCCAGATG 3’ and Reverse 5’ TCCTTATCGATCCCAACAGC 3’. PEMV2 genomic RNA was targeted by forward primer 5’ TTGCAAGGTTCTAGGCATCC 3’ and reverse primer 5’ CAACGATCGAAAAAGACGATG 3’. Gene expression was normalized to the internal control transcripts from the agroinfiltrated p14 RNA silencing suppressor using forward primer 5’ TCCCAAACAGGGGTTTTATG 3’ and reverse primer 5’ GGTAATTGGGAACCCTCGAT 3’. Expression analyses were performed by the ΔΔCq method using Bio-Rad CFX Maestro software. Target fidelity was monitored by melt curve analyses and no reverse transcriptase controls.

